## Supplemental Material for "Ion transfer mechanisms in Mrp-type antiporters from high resolution cryoEM and molecular dynamics simulations"

### Supplementary Materials for

#### High-resolution structure and molecular simulations provide insights into the mechanism of Mrp type antiporters and complex I

Yongchan Lee[1,2,†], Outi Haapanen[3,†], Anton Altmeyer[4,5], Werner Kühlbrandt[1], Vivek Sharma[3,6,*] and Volker Zickermann[4,5,*]

*Corresponding author. Email



**This PDF file includes:**

Supplementary Text
Figs. S1 to S10
Tables S1 to S1-S6
Movie S1
References (65 to 66)

**Other Supplementary Materials for this manuscript include the following:**

Movie S1

**Supplementary text**

**Coordination of internal water molecules by conserved polar and titratable residues**

In MrpA, conserved polar and protonatable residues (Fig. 2, Fig. S3, Fig. S6, Table S4) form a contiguous network extending from the putative proton entry site at Lys408$^{MrpA}$/Glu409$^{MrpA}$ to His248$^{MrpA}$ in the *A* conformation. Water molecules W72 – W78 are arranged around the strictly conserved Lys408$^{MrpA}$. The ligation of the water molecules indicates that Gln309$^{MrpA}$ and Tyr447$^{MrpA}$ and several serine and threonine residues play a crucial role, e.g. water molecule W72 is within hydrogen bonding distance of the highly conserved Ser308$^{MrpA}$ and Tyr447$^{MrpA}$ and moderately conserved Gln309$^{MrpA}$ and Thr444$^{MrpA}$.

From His248$^{MrpA}$ a hydrophilic connection to the cytosolic side is formed by strictly conserved Tyr101$^{MrpA}$, Ser244$^{MrpA}$, Lys299$^{MrpA}$ and highly conserved residues Thr241$^{MrpA}$ and Asp297$^{MrpA}$ (Fig. 2). Tyr101$^{MrpA}$ and Thr241$^{MrpA}$ are within hydrogen bonding distance of W70 and W71. Ser244$^{MrpA}$ binds water W69. Note that residues corresponding to Ser244$^{MrpA}$ also bind a water molecule in respiratory complex I (*21, 22*).

His248 in the *B* conformation is associated with a network of polar residues that connects to the strictly conserved Glu140$^{MrpA}$/Lys223$^{MrpA}$ pair of MrpA. Water W68 is coordinated by the strictly conserved residues Ser146$^{MrpA}$, Thr170$^{MrpA}$ and unconserved Ser147$^{MrpA}$. The adjacent water molecules W66 and W67 are mainly bound by Glu140$^{MrpA}$ and the highly conserved residues Ser143$^{MrpA}$ and Thr222$^{MrpA}$. Interestingly, the indole NH moiety of the strictly conserved Trp139$^{MrpA}$ coordinates both water molecules.

At the interface of MrpA and MrpD, water W65 is bound toTyr136$^{MrpA}$ and the strictly conserved Lys392$^{MrpD}$. In MrpD, we modelled six water molecules (W59-W64) between Lys392$^{MrpD}$ and the highly conserved His332$^{MrpD}$. The water molecules W59 – W64 are further coordinated by Gln307$^{MrpD}$, highly conserved His303$^{MrpD}$ and several polar residues of which Tyr329$^{MrpD}$ is highly conserved. Following the hydrated region towards the core of MrpD, Lys250$^{MrpD}$, His333$^{MrpD}$ and strictly conserved Lys337$^{MrpD}$ form an arrangement that is highly similar to the His349$^{MrpA}$/Lys254$^{MrpA}$/Lys353$^{MrpA}$ triad in MrpA described above. Interestingly, the antiporter from *A. flavithermus* shows a different set of residues in this region of the MrpD subunit. His303$^{MrpD}$, Gln307$^{MrpD}$ and His333$^{MrpD}$ are replaced by Asn303$^{MrpD}$, Ala307$^{MrpD}$ and Asp333$^{MrpD}$, respectively. An extensive analysis of sequence alignments showed that the presence of a Gln or Asn at position 303 is strictly linked with the absence of a Gln at position 307 and the presence of an Asp at position 333 (Fig. S7).

In contrast to MrpA, two water molecules (W56 and W57) are located in the center of MrpD, coordinated by the strictly conserved residues Lys250$^{MrpD}$ and Tyr233$^{MrpD}$ and highly conserved Thr249$^{MrpD}$. A pathway from the center of MrpD to the cytosol is expected but not obvious in the structure. We have proposed that in antiporter-like complex I subunits, conserved residues at the end of TMH10 mark the entrance of a proton channel that can be closed by a conserved phenylalanine residue in TMH11 (*22*). In Mrp, the corresponding residues at the putative channel entry, Asp295$^{MrpD}$ and Lys297$^{MrpD}$, are also conserved and water molecule W58 is bound in close proximity. However, no other water molecules were found between W58 and water molecules in the central axis, which might indicate that the pathway is blocked by the strictly conserved Phe341 in TMH11 (Fig. S3) as recently described for complex I (*22*). An overlay of ND2, ND4 and MrpD shows that the position of this residue agrees well with the "closed" conformation of the ND4 subunit in complex I (Fig. S4).

A hydrated path runs from the center of MrpD to the neighbouring MrpC subunit. Tyr233$^{MrpD}$ bridges between the W56/W57 pair and a cluster of five water molecules (W51 – W55). This cluster is coordinated by Gln166$^{MrpD}$ and the highly conserved Tyr162$^{MrpD}$. The latter residue engages in a hydrogen bond with the strictly conserved Ser143$^{MrpD}$. Water molecule W50 is bound between Lys219$^{MrpD}$ and Glu137$^{MrpD}$, the strictly conserved Lys/Glu pair of MrpD, and is further ligated by Ser170$^{MrpD}$. We modelled Lys219$^{MrpD}$ in two conformations, only one of which allows for a hydrogen bond to the water molecule.

In the neighbouring MrpC subunit, TMH2 and 3 are at the center of a highly hydrated area. A cluster of water molecules W32 – W37 at the interface of MrpC and MrpD is coordinated by Glu137$^{MrpD}$ and by moderately conserved Ser36$^{MrpC}$, His40$^{MrpC}$ (see below), Ser80$^{MrpC}$, Thr84$^{MrpC}$. Towards the C-terminal domain of MrpA, a large water (W21 – W31) is bound by polar residues of MrpC, His37$^{MrpC}$ and His40$^{MrpC}$, as well as Thr690$^{MrpA}$ and strictly conserved Gln683$^{MrpA}$ and Glu687$^{MrpA}$. A polar residue at position 690 of MrpA is strictly conserved. Residues His37$^{MrpC}$ and His40$^{MrpC}$ in TMH2 were recently described as being critical for sodium binding (*14*). We note that neither residue is strictly conserved, but histidine or asparagine are the only residues allowed at position 40. A string of water molecules W16 – W20 connects TMH18 of MrpA with the highly conserved Asp38$^{MrpF}$. Residue Ser75$^{MrpF}$ was modelled in two conformations one of which binds water W16. Two clusters of waters (W6 – W15) are located at the interface of MrpF and MrpG. The larger cluster is arranged around Asp38$^{MrpF}$ and further binding interactions exist with the adjacent Thr39$^{MrpF}$, Ser68$^{MrpF}$ and Thr40$^{MrpG}$, Thr44$^{MrpG}$ and Thr79$^{MrpG}$. Polar residues at positions 40, 44 and 79 of MrpG are highly conserved. The smaller cluster is coordinated by residues from both subunits but none of them is conserved.

A previous study suggested that sodium entry from the cytoplasm occurs at the interface between MrpG and MrpE (*14*). We find water molecules W1-W5 distributed around the proposed sodium entry site and several of the hydrogen-bonding residues including highly conserved His37$^{MrpG}$, Thr116$^{MrpE}$, and strictly conserved Thr113$^{MrpE}$, His131$^{MrpE}$. A sodium exit (*14*) or proton entry (*18*) site was proposed near the highly conserved residues Asp776$^{MrpA}$ and Glu780$^{MrpA}$ in TMH21 of MrpA. Water molecules W45 and W46 are close to the critical residues and are further coordinated by strictly conserved residues Thr777$^{MrpA}$ and Thr75$^{MrpC}$. Water molecules W38 – W44 are found in a cavity that may form a pathway to the cytoplasm. They are coordinated by the strictly conserved residues Asp678$^{MrpA}$, Asn766$^{MrpA}$, and Asp121$^{MrpB}$. Asn766$^{MrpA}$ was modelled in two different conformations oriented towards different waters in the hydrated path.

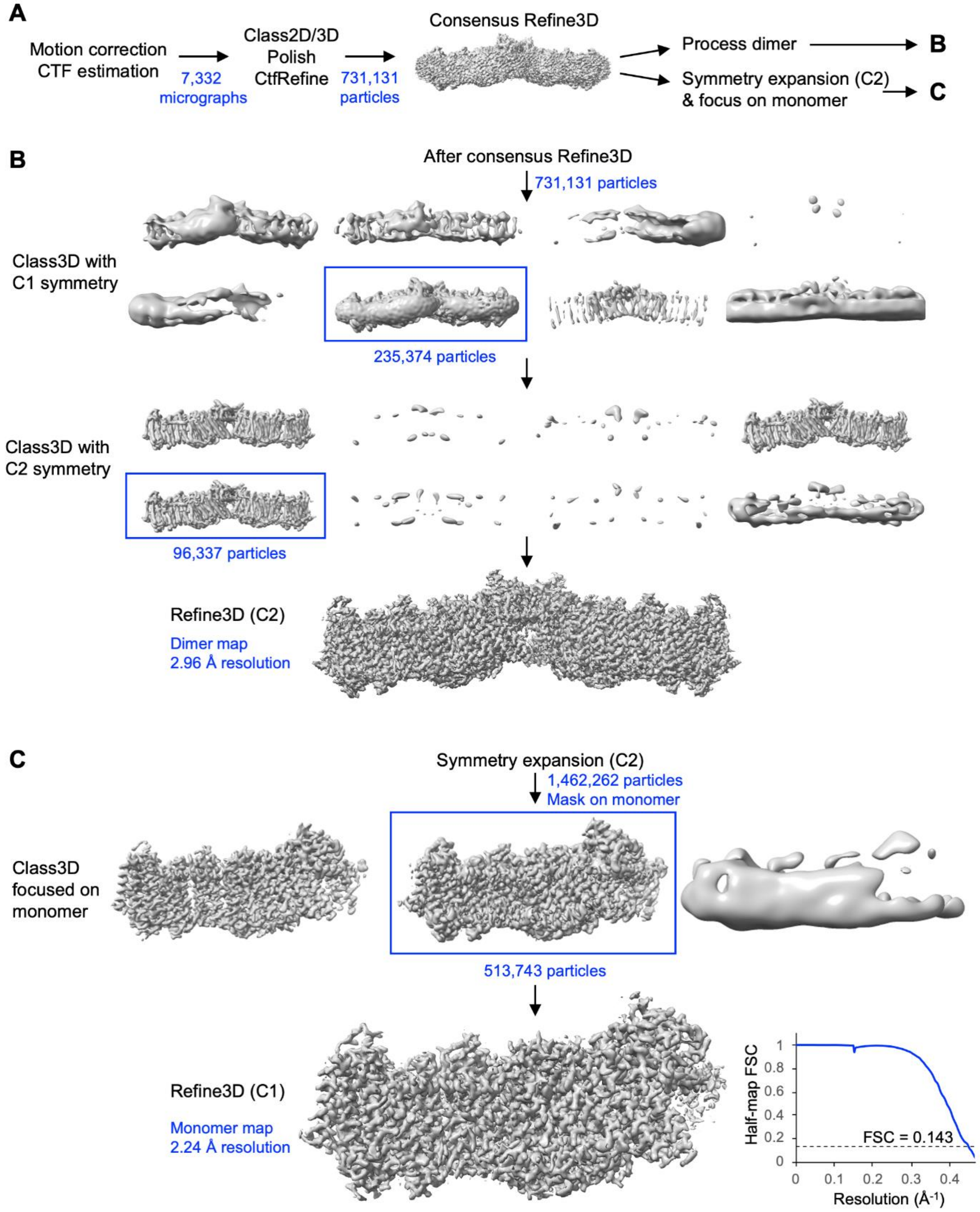


**Figure S1.**

**Workflow of single-particle data processing for *B. pseudofirmus* Mrp. (A)** Initial stages of data processing, before particles were subjected to different processing strategies. **(B)** Dimer refinement. **(C)** Monomer refinement.

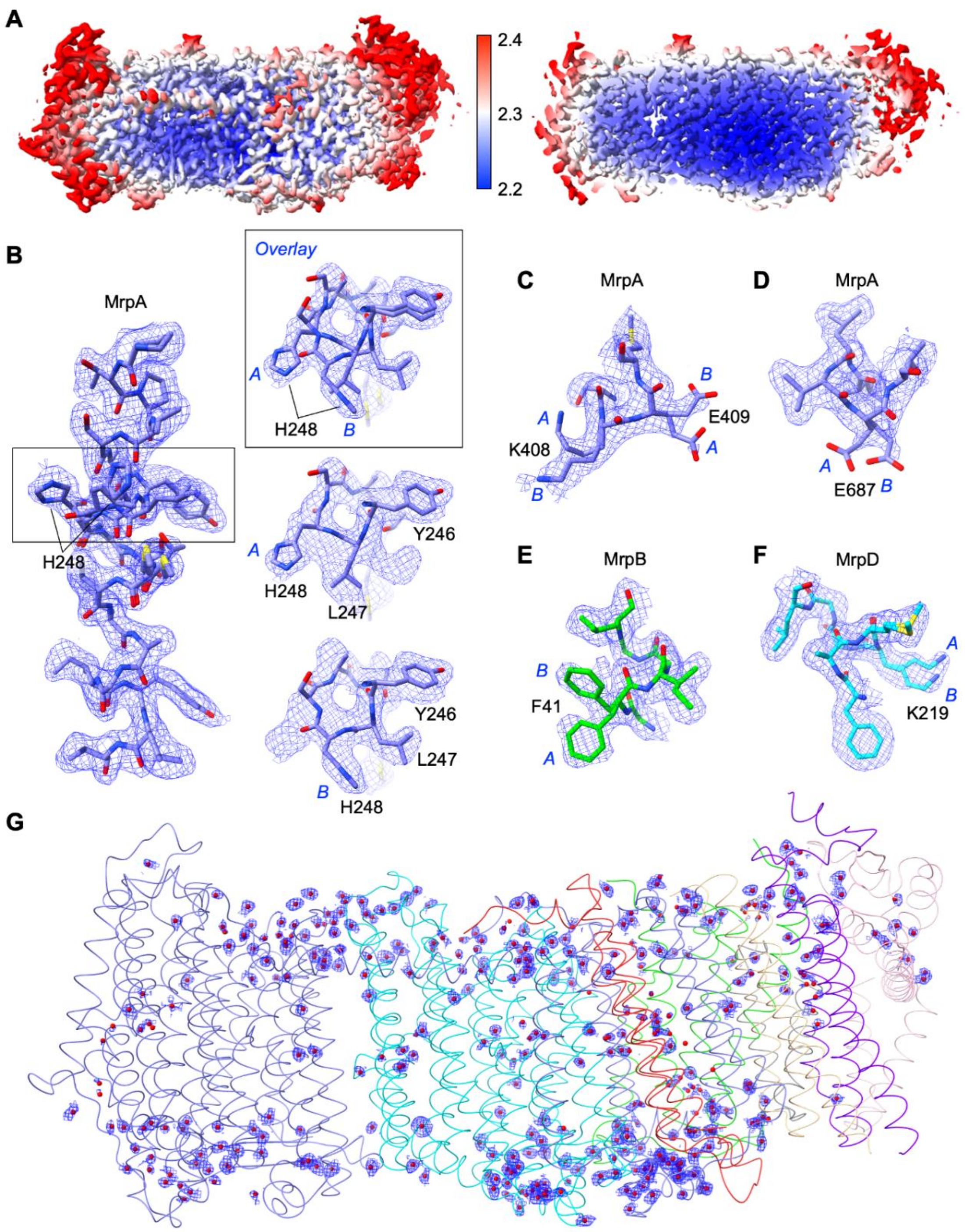


**Figure S2.**

**Local resolution and examples of cryo-EM densities. (A)** Local resolution map of the *B. pseudofirmus* Mrp monomer**. (B)** Two alternative conformations of residues 246 – 252 in MrpA. Overlaid models from two different views are outlined. Separated models of two conformations are shown below. **(C)** Alternative sidechain conformations of K408 and E409 in MrpA. **(D)** Alternative sidechain conformations of E687 in MrpA. **(E)** Alternative sidechain conformations of F41 in MrpB. **(F)** Alternative sidechain conformations of K219 in MrpD**. (G)** All solvent densities that were modelled as water.

A MrpA-ND5:

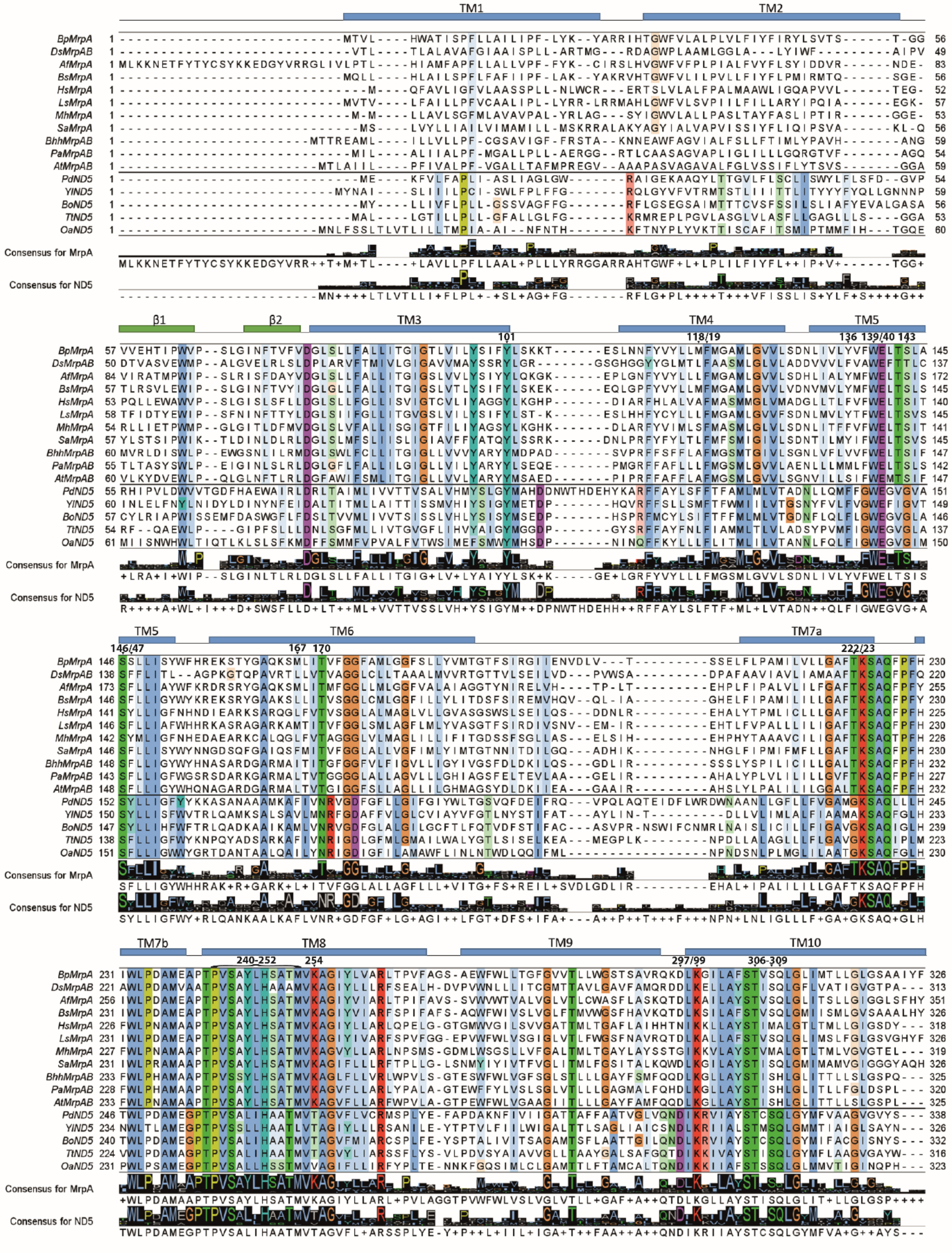

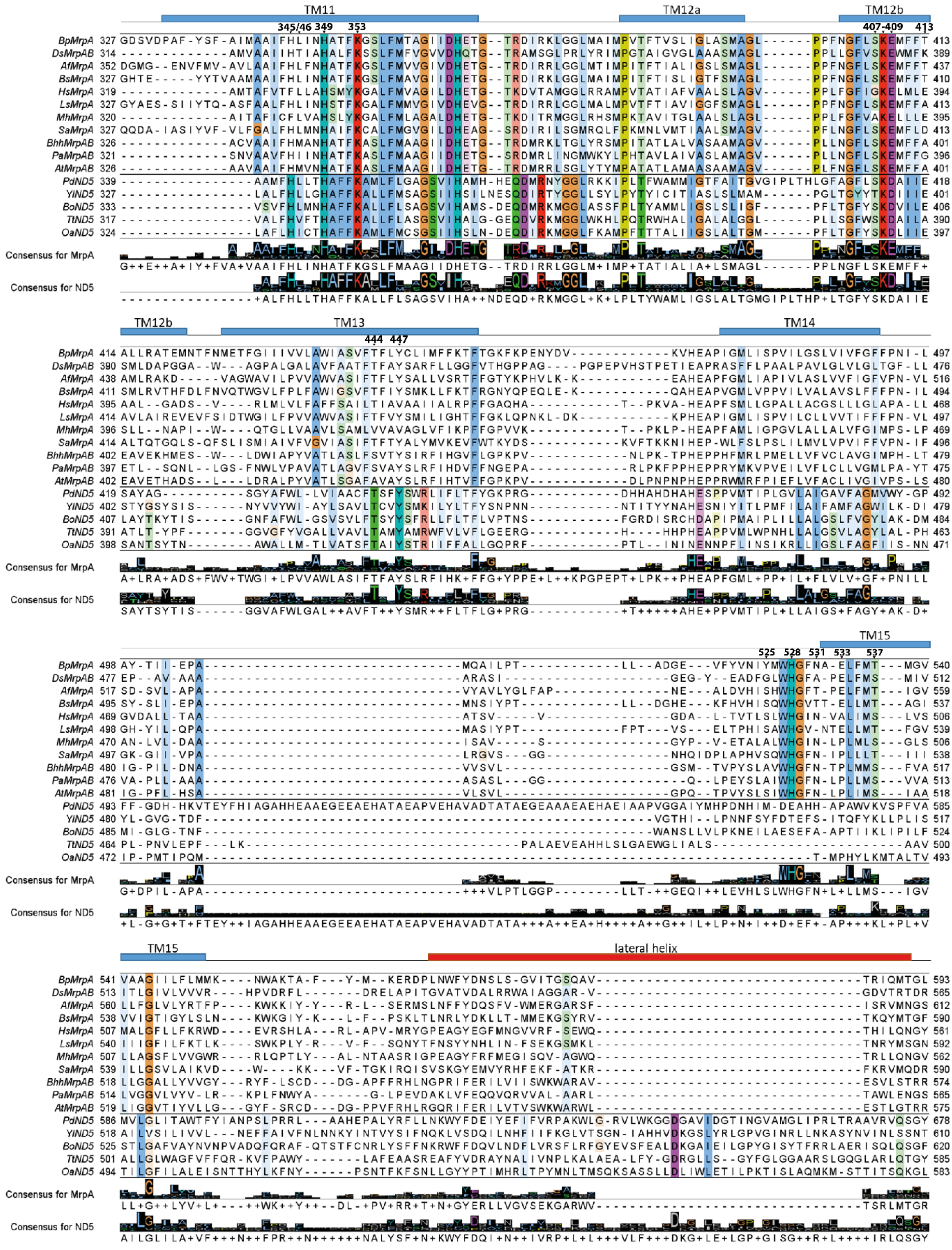

TM11
TM12a
TM12b
345/46 349 353
407-409 413
BpMrpA
DsMrpAB
AfMrpA
BsMrpA
HsMrpA
LsMrpA
MhMrpA
SaMrpA
BhhMrpAB
PaMrpAB
AtMrpAB
PdND5
YlND5
BoND5
TtND5
OaND5
Consensus for MrpA
Consensus for ND5
TM12b
TM13
TM14
444 447
TM15
525 528 531 533 537
TM15
lateral helix

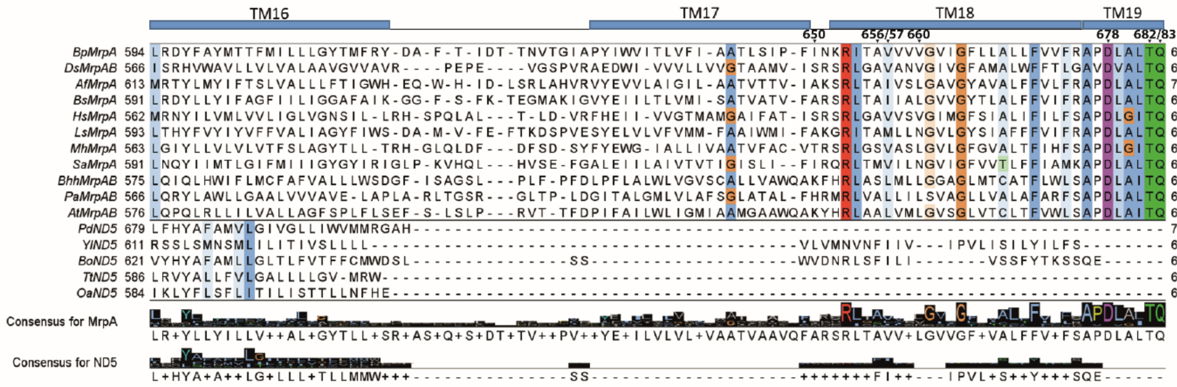
TM16
TM17
TM18
TM19
BpMrpA
DsMrpAB
AfMrpA
BsMrpA
HsMrpA
LsMrpA
MhMrpA
SaMrpA
BhhMrpAB
PaMrpAB
AtMrpAB
PdND5
YlND5
BoND5
TtND5
OaND5
Consensus for MrpA
Consensus for ND5

B MrpA-ND6:

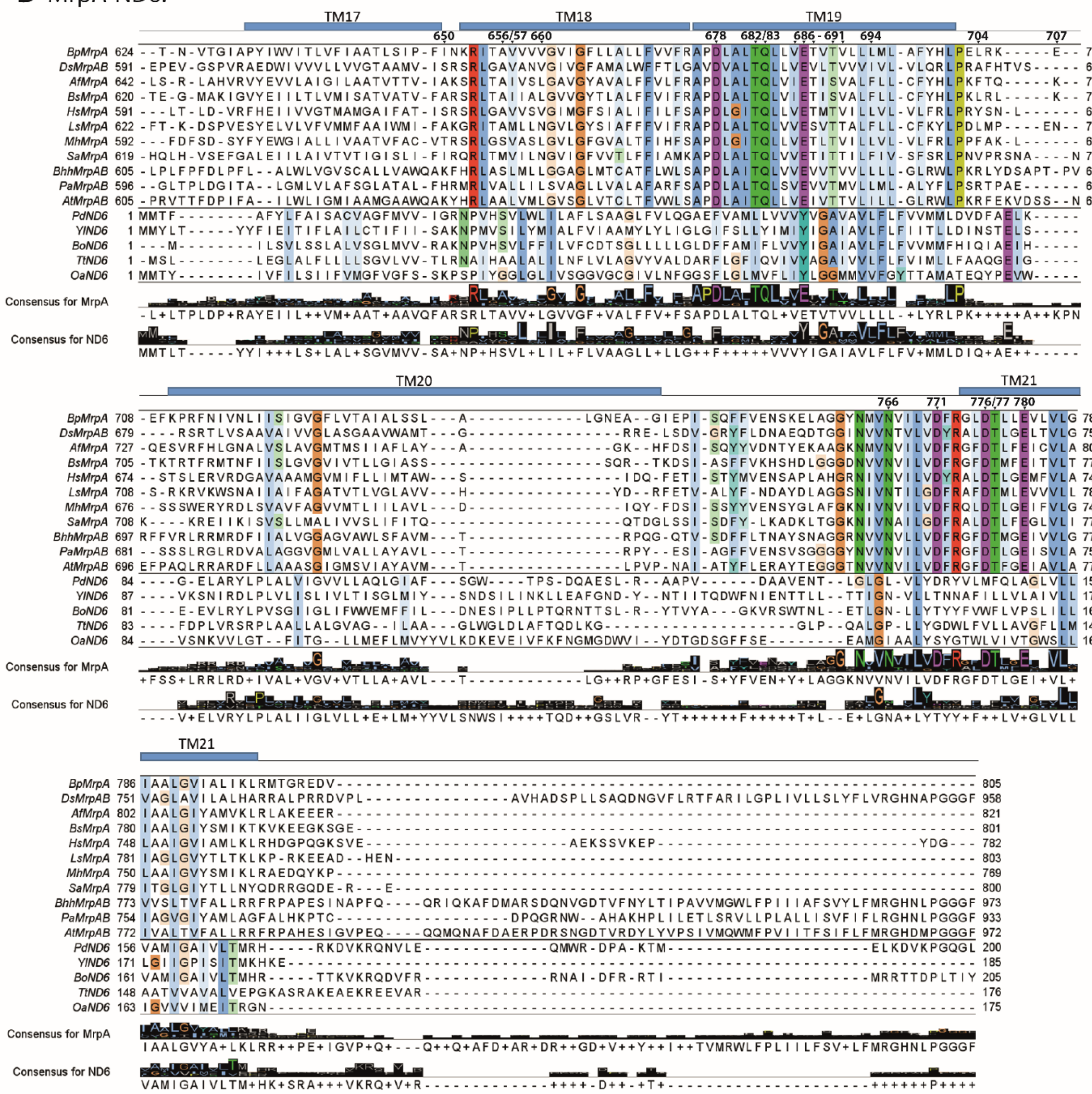
TM17
TM18
TM19
TM20
TM21
BpMrpA
DsMrpAB
AfMrpA
BsMrpA
HsMrpA
LsMrpA
MhMrpA
SaMrpA
BhhMrpAB
PaMrpAB
AtMrpAB
PdND6
YlND6
BoND6
TtND6
OaND6
Consensus for MrpA
Consensus for ND6

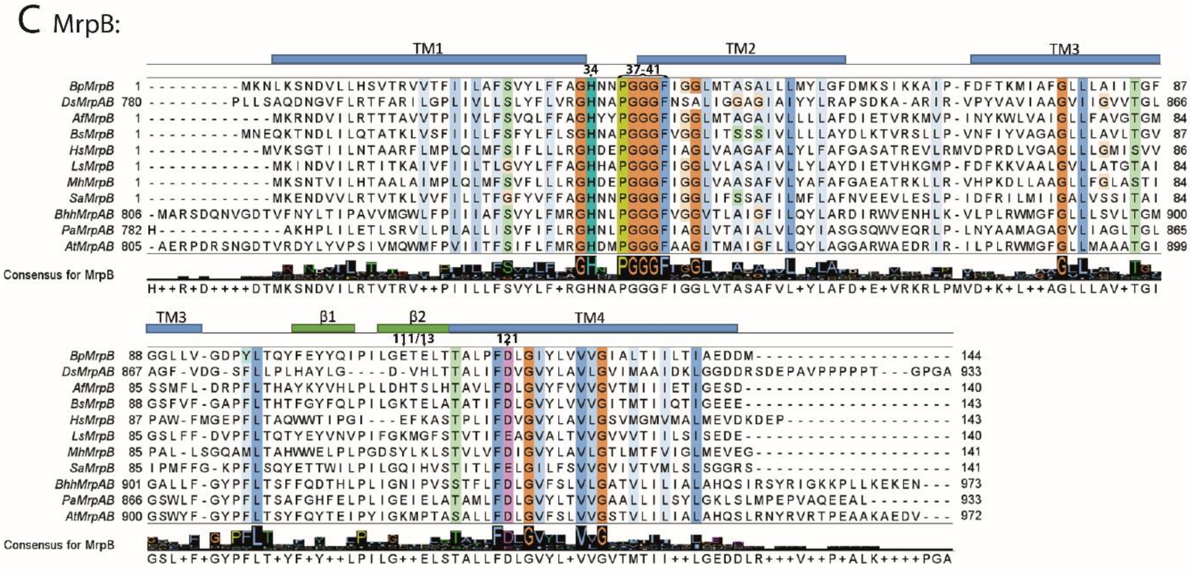
C MrpB:
TM1
TM2
TM3
TM3
β1
β2
TM4
BpMrpB
DsMrpAB
AfMrpB
BsMrpB
HsMrpB
LsMrpB
MhMrpB
SaMrpB
BhhMrpAB
PaMrpAB
AtMrpAB
Consensus for MrpB

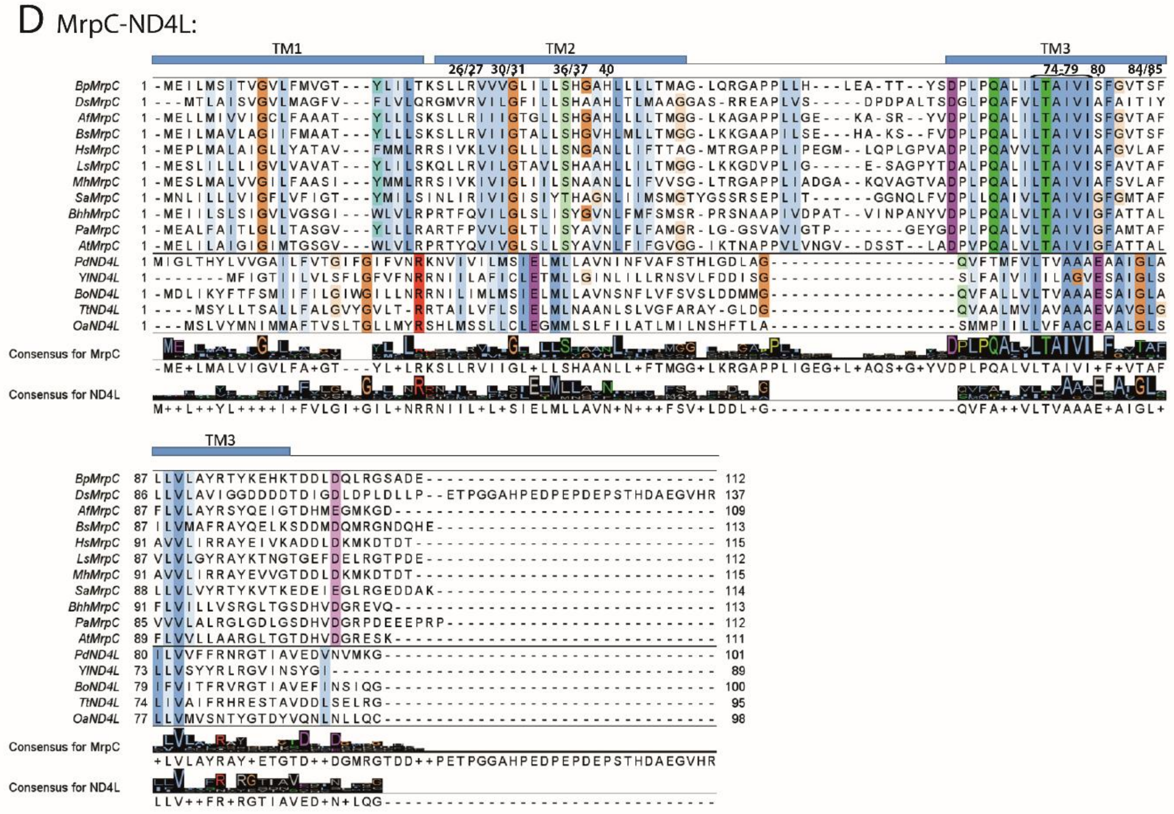
D MrpC-ND4L:
TM1
TM2
TM3
TM3
BpMrpC
DsMrpC
AfMrpC
BsMrpC
HsMrpC
LsMrpC
MhMrpC
SaMrpC
BhhMrpC
PaMrpC
AtMrpC
PdND4L
YlND4L
BoND4L
TtND4L
OaND4L
Consensus for MrpC
Consensus for ND4L

### E MrpD-ND2/ND4:

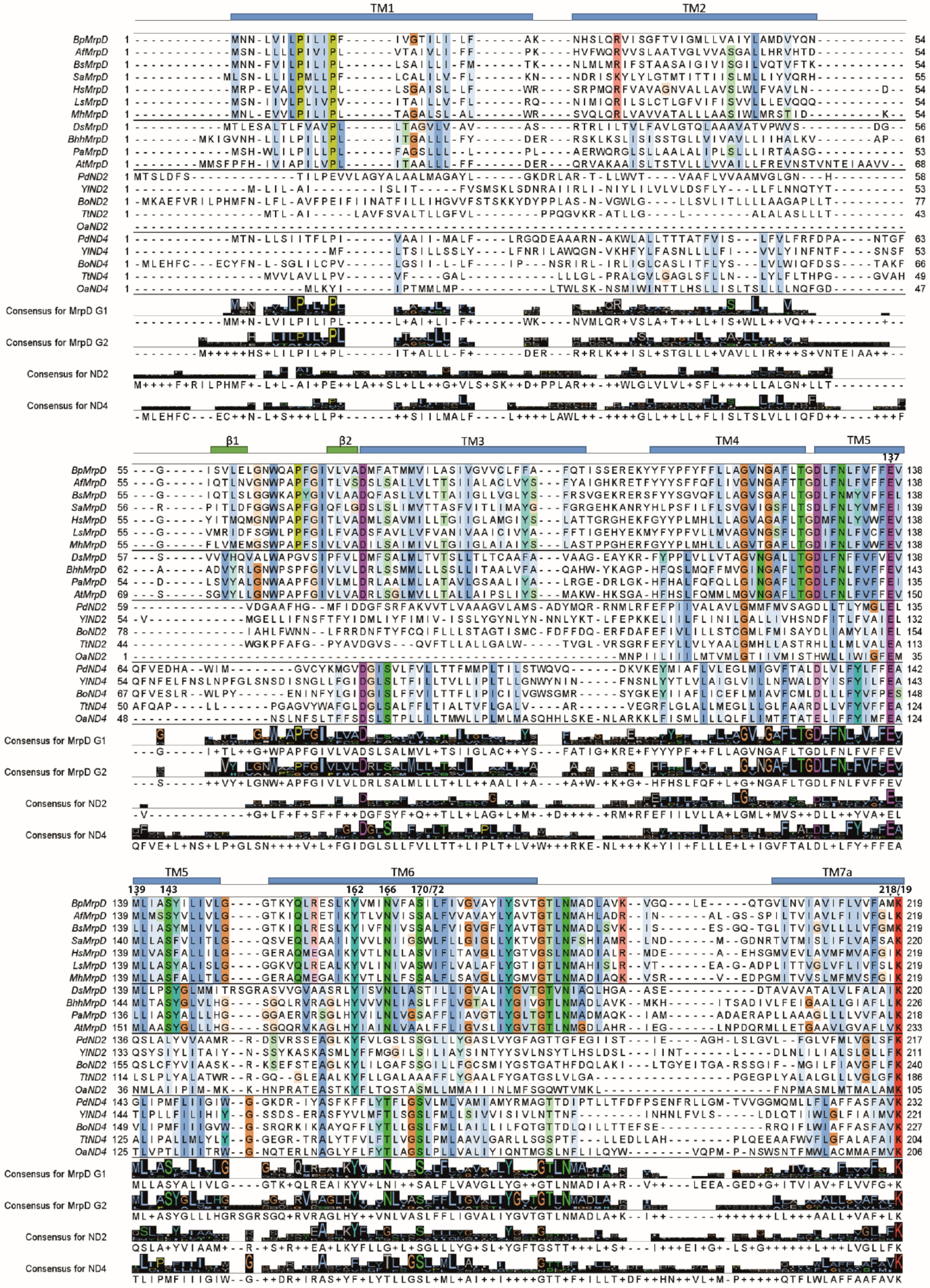

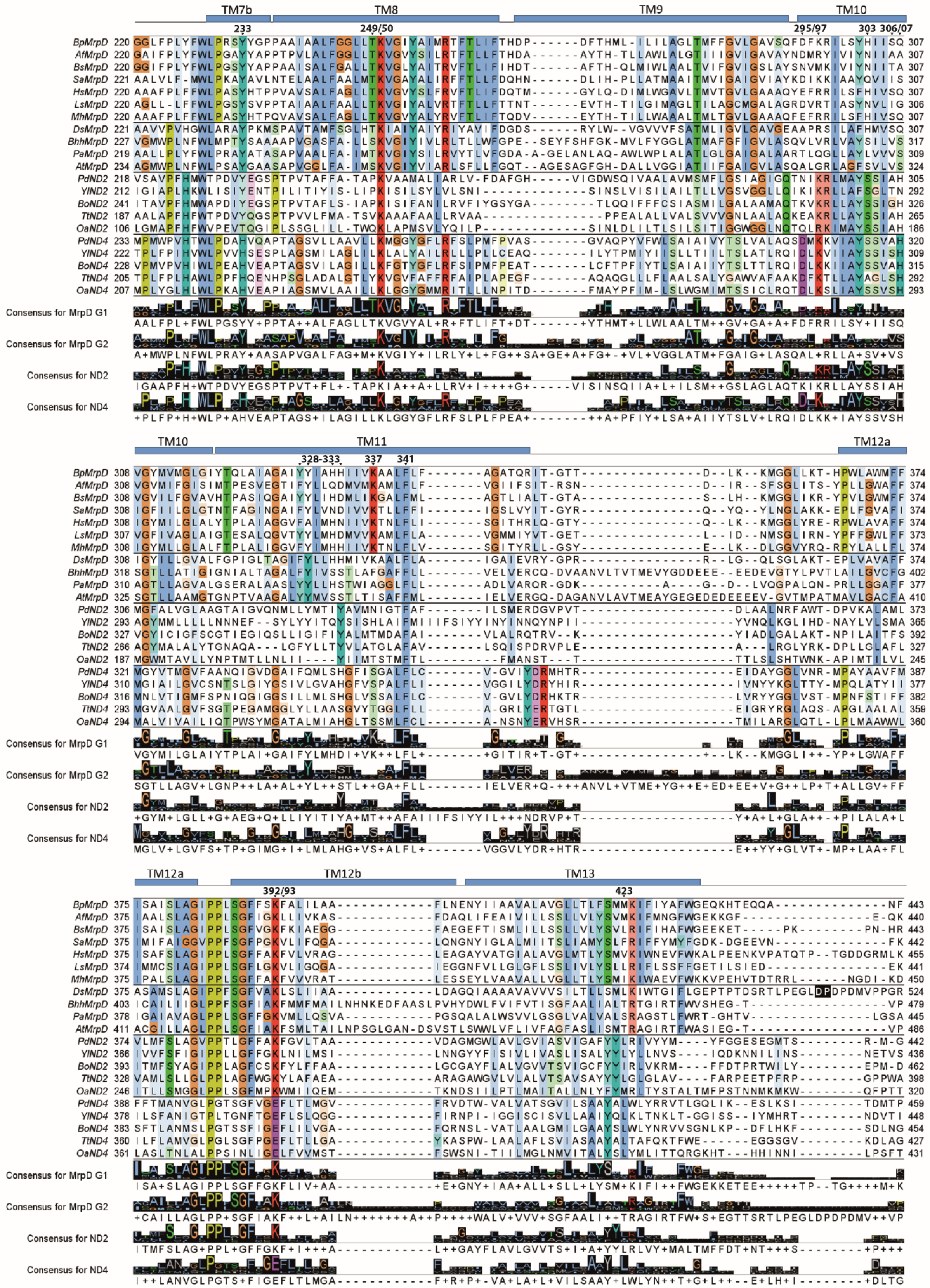
TM7b
TM8
TM9
TM10
233
249/50
295/97
303 306/07
BpMrpD
AfMrpD
BsMrpD
SaMrpD
HsMrpD
LsMrpD
MhMrpD
DsMrpD
BhhMrpD
PaMrpD
AtMrpD
PdND2
YlND2
BoND2
TtND2
OaND2
PdND4
YlND4
BoND4
TtND4
OaND4
Consensus for MrpD G1
Consensus for MrpD G2
Consensus for ND2
Consensus for ND4
TM11
TM12a
328-333
337
341
TM12b
TM13
392/93
423

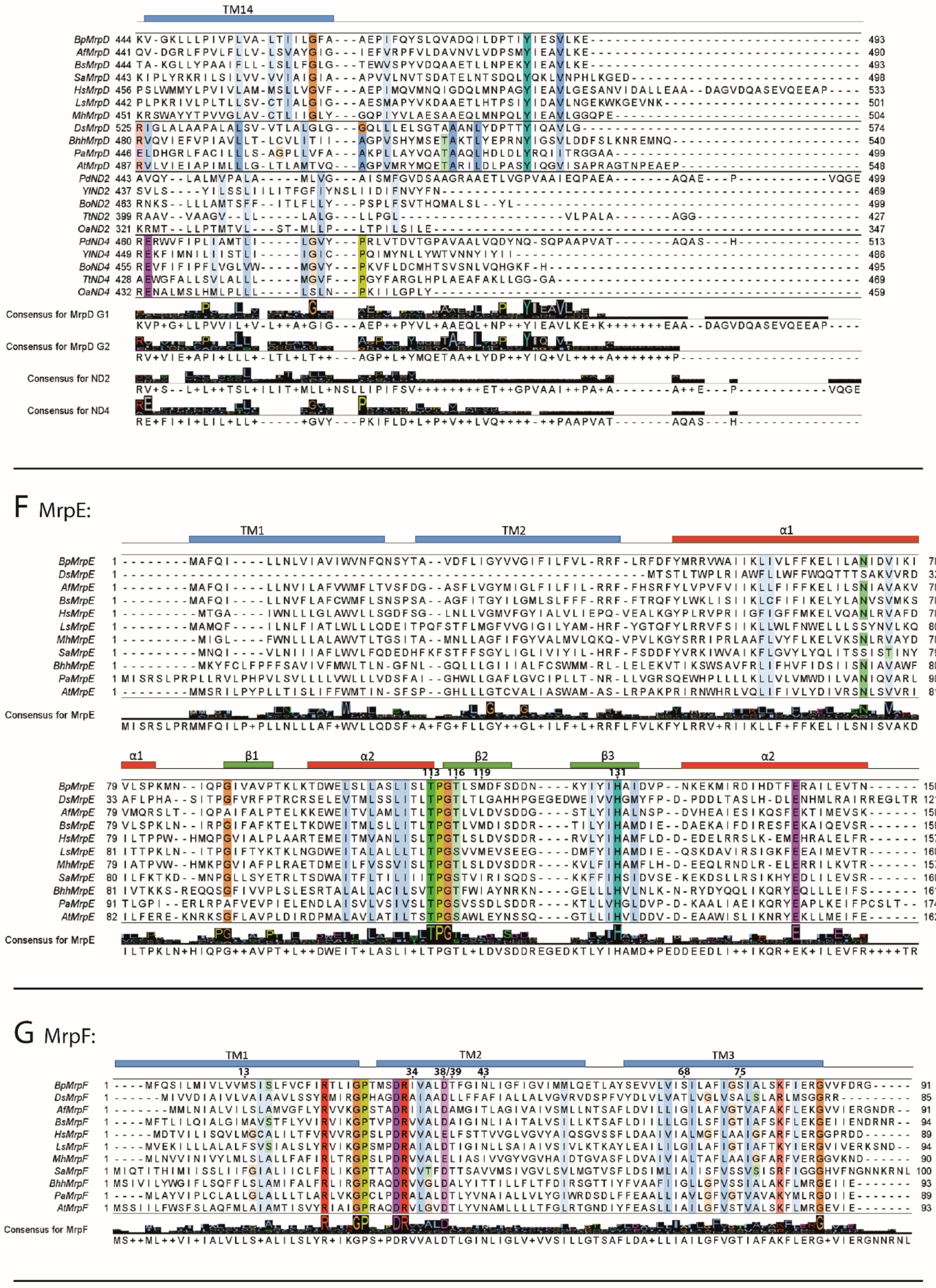
TM14
Consensus for MrpD G1
Consensus for MrpD G2
Consensus for ND2
Consensus for ND4
F MrpE:
TM1
TM2
α1
β1
α2
β2
β3
Consensus for MrpE
G MrpF:
TM1
TM2
TM3
Consensus for MrpF

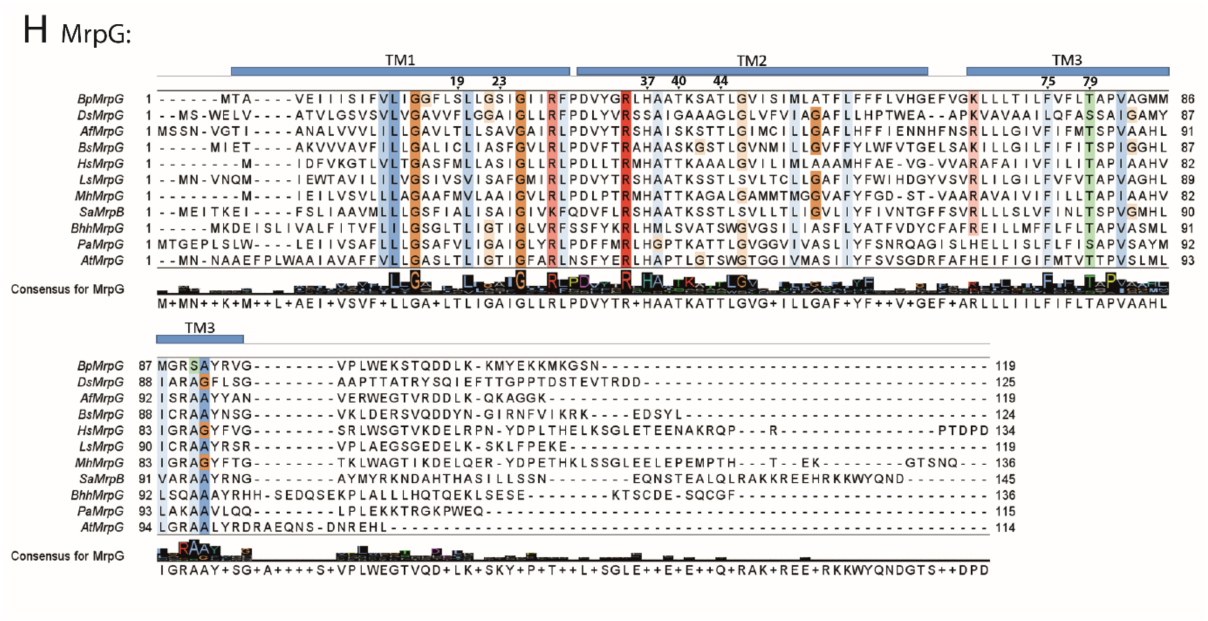


**Figure S3.**

**Sequence alignment of orthologous subunits in Mrp antiporters and respiratory complex I.** Alignments include the Mrp antiporter subunits A-G and A'-G from the species *Bacillus pseudofirmus* (BpMrp), *Anoxybacillus flavithermus* (AfMrp), *Bacillus subtilis* (BsMrp), *Staphylococcus aureus* (SaMrp), *Halomonas* sp. (HsMrp), *Lysinibacillus sphaericus* (LsMrp), *Marinobacter hydrocarbonoclasticus* (MhMrp), *Dietzia* sp. (DsMrp), *Bartonella henselae* (BhhMrp), *Pseudomonas aeruginosa* (PaMrp) and *Agrobacterium tumefaciens* (AtMrp), as well as the complex I subunits ND2, ND4, ND5, ND6 and ND4L from the species *Paracoccus denitrificans* (PdND), *Yarrowia lipolytica* (YlND), *Brassica oleracea* (BoND), *Thermus thermophilus* (TtND) and *Ovis aries* (OaND).

The alignments were created using ClustalO (*36*) with default values in Jalview 2.11.1.4 (*35*) and are color-coded in the ClustalX (*38*) preset filtered by 50 % conservation. Selected residues are marked above the alignments with their sequence position in BpMrp. The secondary structure elements of BpMrp subunits are indicated as blue (transmembrane helices: TM), red (helices outside the membrane: α) or green (β-sheets: β) boxes. **(A)** Alignment of subunits MrpA, MrpA' and ND5. **(B)** Alignment of the subunits MrpA, MrpA' and ND6. **(C)** Alignment of subunit MrpB. **(D)** Alignment of subunits MrpC and ND4L. **(E)** Alignment of subunits MrpD, ND2 and ND4. The MrpD sequences are differentiated between group 1 (separate MrpA and MrpB) and group 2 (MrpA' fusion protein) operons. A 68 residue insert between Asp447 and Pro515 in DsMrpD is not displayed (position highlighted in black). **(F)** Alignment of subunit MrpE. **(G)** Alignment of subunit MrpF. **(H)** Alignment of subunit MrpG.

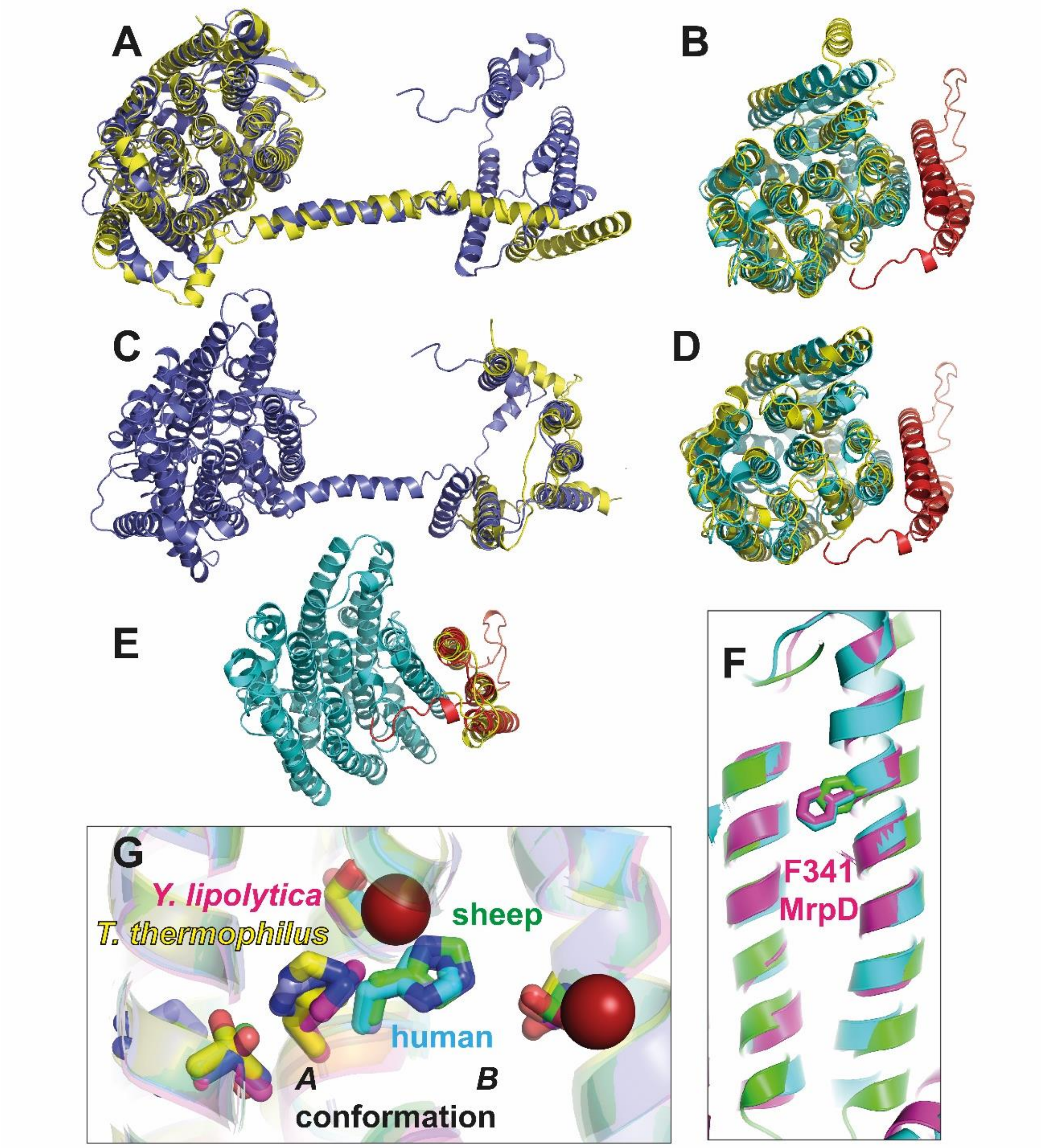


**Figure S4.**

**Structural features of the Mrp antiporter conserved in complex I.** Overlay of complex I (*Y. lipolytica*, PDB 7O71, yellow) and Mrp antiporter subunits (color as in Fig. 1), **(A)** MrpA/ND5, **(B)** MrpD/ND4, **(C)** MrpA/ND6, **(D)** MrpD/ND2, **(E)** MrpC/ND4L. **(F)** The position of a highly conserved phenylalanine residue opens or closes a proton pathway in subunits ND2 (green, open) and ND4 (cyan, closed); the position of the corresponding residue Phe341 in MrpD (magenta) matches the closed conformation. **(G)** In complex I structures, residues corresponding to His248$^{MrpA}$ are either in the *A* (e.g. *T. thermophilus* PDB 6I1P, *Y. lipolytica* PDB 7O71) or *B* conformation (e.g. human PDB 5XTD, sheep PDB 6ZKA); residues corresponding to Ser146$^{MrpA}$, Ser244$^{MrpA}$, and Thr306$^{MrpA}$ (compare Fig. 2) are strictly conserved in complex I and form connections to three pathways for protons.

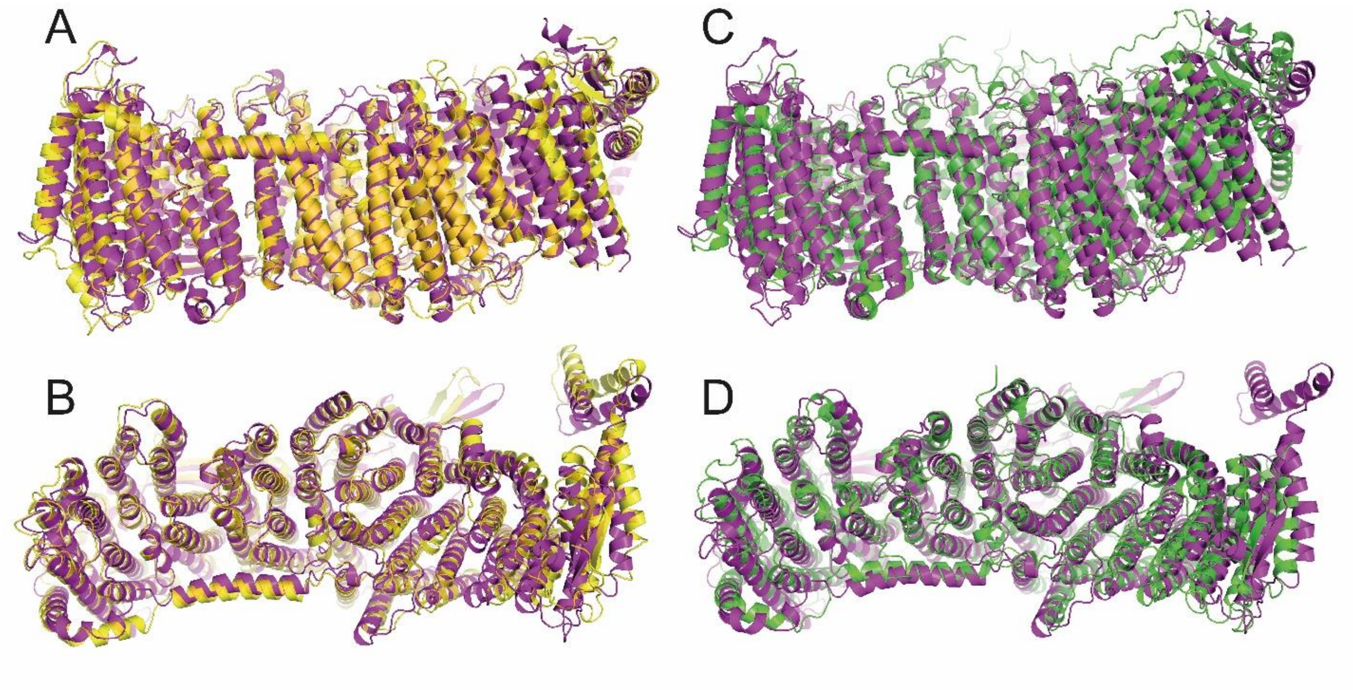


**Figure S5.**

**Mrp antiporter from *B. pseudofirmus* compared with Mrp antiporters from *A. flavithermus* and *Dietzia* sp. (A)** Side view and **(B)** top view of an overlay of Mrp antiporters from *B. pseudofirmus* (magenta) and *A. flavithermus* (yellow, PDB ID 6Z16); note that both structures show one protomer of a dimeric complex. **(C)** Side view and **(D)** top view of an overlay of Mrp antiporters from *B. pseudofirmus* (magenta) and *Dietzia sp.* (green, PDB ID 7D3U); note that the Mrp antiporter from *Dietzia sp.* was isolated as a monomeric complex and that MrpA and MrpB are fused.

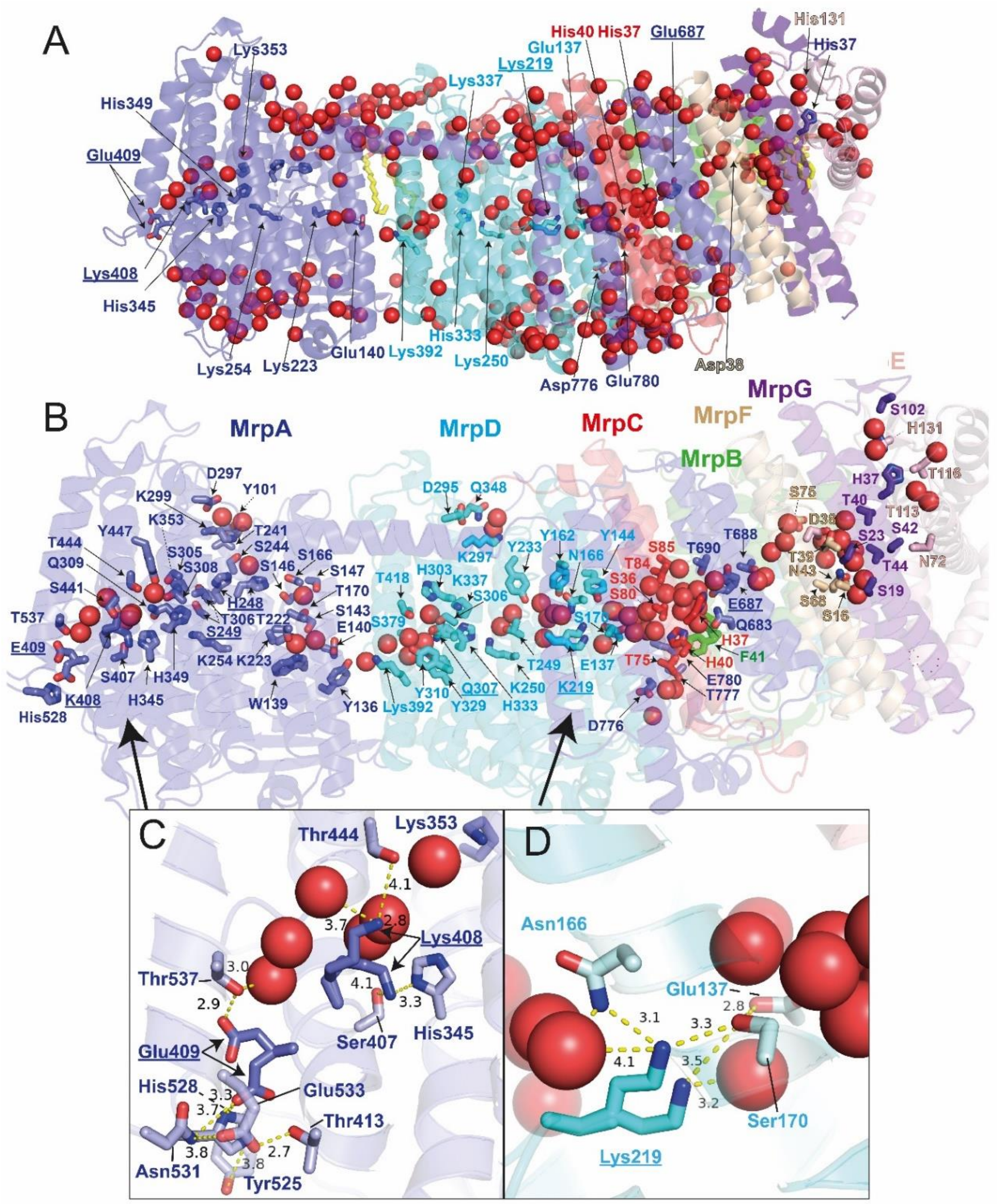


**Figure S6.**

**Water molecules and liganding residues in the *B. pseudofirmus* Mrp antiporter. (A)** A total of 360 water molecules (red spheres) were modelled. More than 70 water molecules define putative ion translocation pathways in the complex interior (compare Fig. 2). **(B)** Detailed view including polar residues not shown in Fig. 2. **(C)** Glu409$^{MrpA}$ interacts with strictly conserved His528$^{MrpA}$ and engages in a hydrogen bonding network at the periplasmic proton entry site; in a second conformation it points towards Thr537$^{MrpA}$ and the water chain leading to His248$^{MrpA}$. Lys408$^{MrpA}$ changes between a position pointing towards highly conserved Ser407 and His345 and a position where it binds two water molecules. **(D)** Lys219$^{MrpD}$ either orients towards waters in the center of MrpD or to a water molecule that is liganded by Ser170$^{MrpD}$ and Glu137$^{MrpD}$ of the MrpD K/E pair.

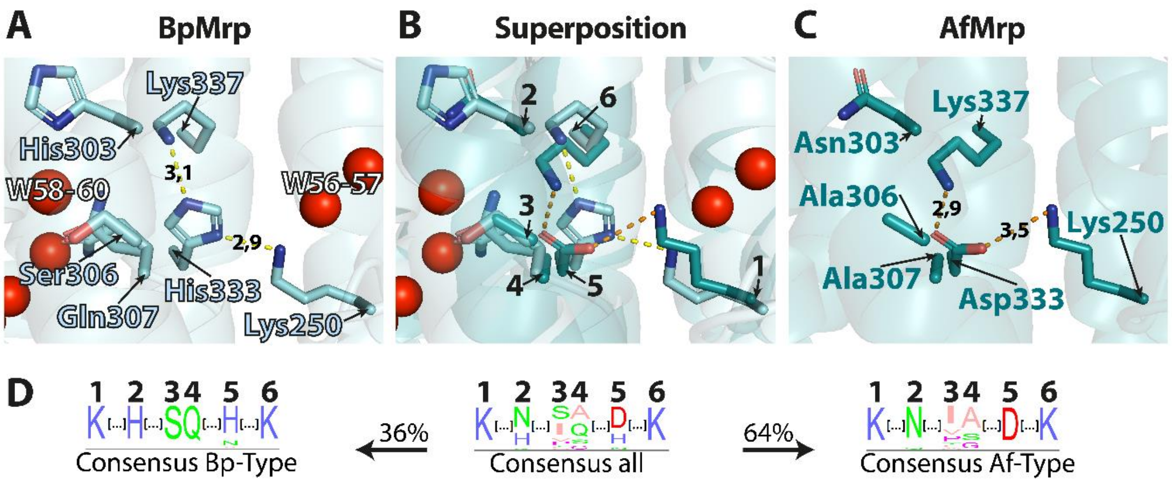


**Figure S7.**

**Co-evolution of residues in MrpD of Group 1 Mrp antiporter operons**. **(A)** Close-up view of the region around the strictly conserved Lys250 in TMH8 of MrpD of *B. pseudofirmus* (BpMrp) (compare Fig. 2). **(B)** Superposition of (A) and (C). **(C)** Close-up view on the region around the strictly conserved Lys250 in TMH8 of MrpD of *Anoxybacillus flavithermus* (AfMrp; PDB: 6Z16). **(D)** Consensus of 1200 sequences of Mrp group 1 operons for the residues shown in (B). Two clear subgroups can be identified. The Bp-Type (present in about 1/3 of the sequences) expresses His in position 2, Ser in 3, Gln in 4 and His or Asn in 5. The Af-Type (present in about 2/3 of the sequences) expresses Asn or Gln in position 2, varying residues in 3, small residues in 4 and Asp in 5.

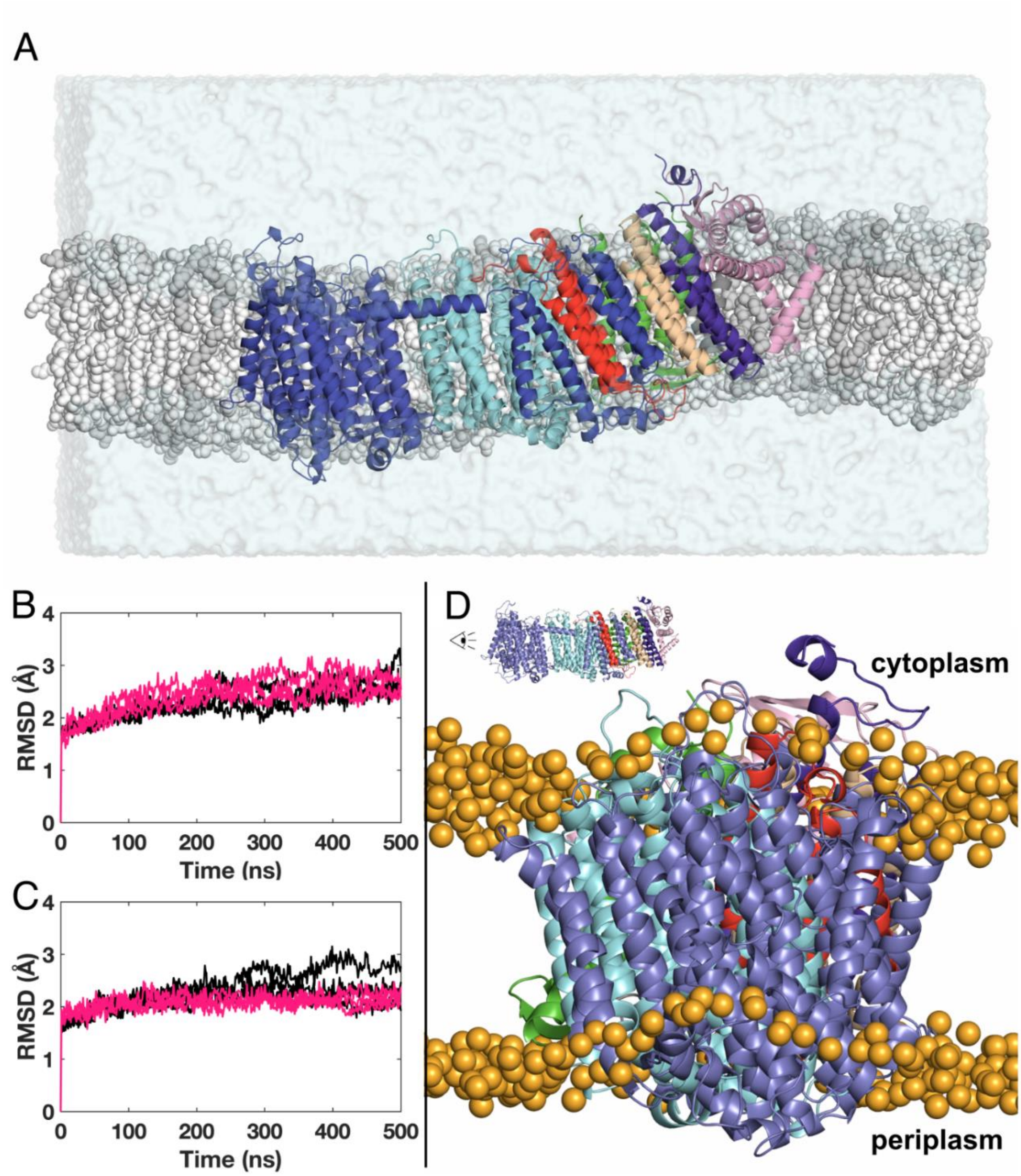


**Figure S8.**

**MD simulation model systems and lipid bilayer arrangement near the proton entry site in MrpA.**
**(A)** Model system of Mrp antiporter (cartoon representation) embedded in a hybrid lipid membrane (grey spheres) and solvent (turquoise surface representation, $Na^+$ and $Cl^-$ ions omitted for clarity). POPE lipids are shown as white and POPG as grey spheres. Protein is colored as in Figure 1. **(B)** RMSD of protein CA atoms in SA1 (black) and SB1 (pink) simulations. **(C)** RMSD of protein CA atoms in PA1 (black) and PB1 (pink) simulations. **(D)** Simulation snapshot reveals bending in the lipid membrane occurs closer to the periplasmic surface of the putative proton-uptake site. Protein is shown in ribbons representation colored as in Figure 1. Phosphorus atoms of lipids surrounding the protein are shown as yellow spheres. Lipid fatty acid side chains are omitted for clarity.

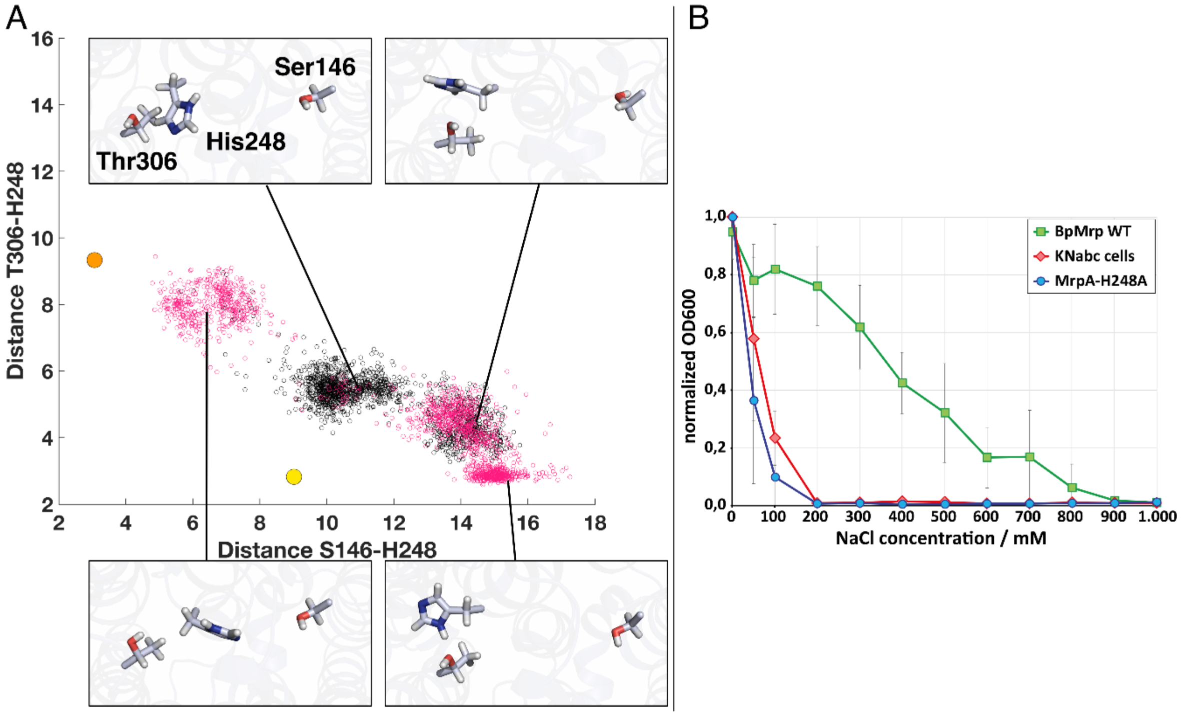


**Figure S9.**

**His248$^{MrpA}$ is a critical residue**. **(A)** Variation in sidechain of neutral His248$^{MrpA}$ (with δ nitrogen protonated) in S state simulations where all amino acids are in their standard states. The scatter plot shows distances between NE2 atom of His248$^{MrpA}$ and OG atoms of Ser146/Thr306$^{MrpA}$ from SA (pink dots) or SB (black dots) simulations. Solid yellow and orange spheres indicate the structural distances seen in alternative *A* and *B* conformations. Note that in the S state simulations, conformations of His248$^{MrpA}$ are not populated (no overlap with solid orange and yellow circles), whereas they are clearly observed in P state simulations, where protonation states of sidechains are determined by pKa calculations (Fig. 5A). **(B)** Complementation of *E. coli* strain KNabc without intrinsic antiport activity (red curve) with an expression vector carrying the wild-type Mrp operon of *B. pseudofirmus* sustains salt tolerant growth (green curve) in contrast to complementation with the His248$^{MrpA}$ mutant (blue curve).

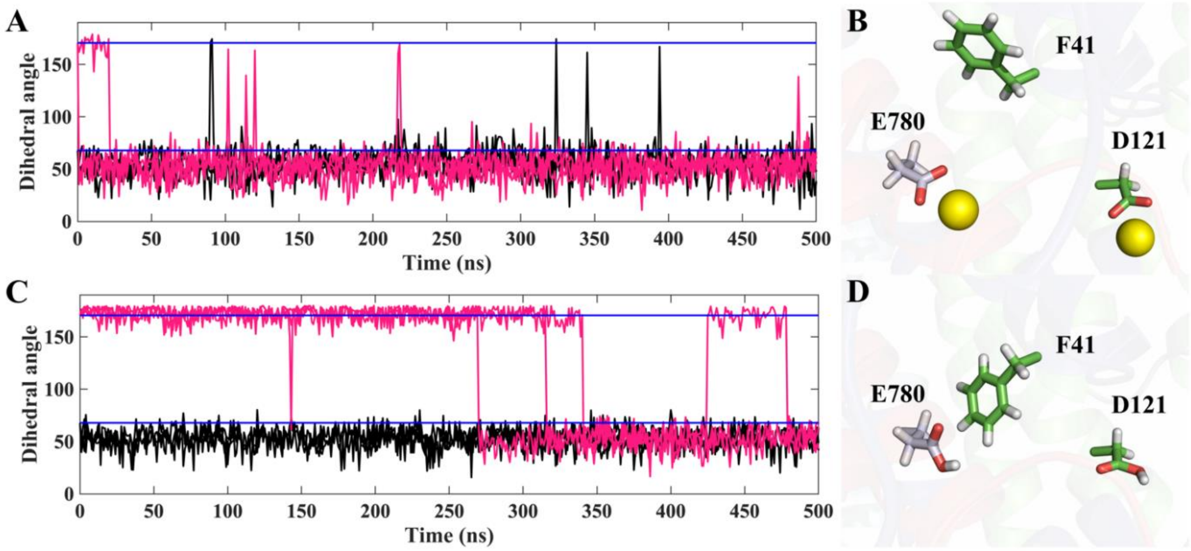


**Figure S10.**
**Phe41$^{MrpB}$ dynamics and sodium gating. (A)** Dihedral angle (C-CA-CB-CD) of Phe41$^{MrpB}$ from SA1 (3 x black) and SB1 (3 x pink) simulations. **(B)** Snapshot at 200 ns from SA1 simulation. Carbon atoms in the MrpB subunit are green; in MrpA they are lilac. In the simulations, two sodium atoms bind to residues in their charged residues spontaneously. Phe41$^{MrpB}$ adopts the *A* conformation regardless of the initial structure as is also seen in panel (A). **(C)** Dihedral angle (C-CA-CB-CD) of Phe41$^{MrpB}$ from PA1 (3 x black) and PB1 (3 x pink) simulations. **(D)** Snapshot at 200 ns from one PB1 simulation. Colors as in (B). Since Glu780$^{MrpA}$ and Asp121$^{MrpB}$ are uncharged in the PB1 simulation, no sodium ions interact with them, allowing Phe41 to remain in the *B* conformation.

**Table S1.**

**Cryo-EM data and model statistics**

| | *B. pseudofirmus* Mrp | |
|---|---|---|
| **Data collection** | | |
| Microscope | Titan Krios 2 | |
| Camera | K3 | |
| Magnification | 105,000 | |
| Voltage (kV) | 300 | |
| Electron exposure ($e^-/Å^2$) | 50.0 | |
| Defocus range (µm) | −0.8 to −2.0 | |
| Calibrated pixel size (Å) | 0.837 | |
| | Monomer | Dimer |
| **Data processing** | | |
| Final particle images (no.) | 513,743 | 96,337 |
| Final pixel size (Å) | 1.07136 | 1.07136 |
| Symmetry imposed | C1 | C2 |
| Map resolution (Å) | | |
| Half map FSC = 0.143 | 2.24 | 2.96 |
| Map sharpening *B* factor ($Å^2$) | –16.7 | –34.5 |
| Local resolution range (Å) | 2.2–2.8 | |
| **Refinement** | | |
| Initial model (PDB codes) | 6Z16 | |
| Refinement resolution (Å) | 2.20 | |
| Model resolution (Å) | | |
| Map-model FSC = 0.5 | 2.24 | |
| Model composition | | |
| Non-hydrogen atoms | 16,100 | |
| Protein residues | 1,957 | |
| Water | 360 | |
| Other ligands | 3 (POPE) | |
| Average *B* factors ($Å^2$) | | |
| Protein | 48.3 | |
| Water | 50.7 | |
| Other ligands | 109.7 | |
| R.m.s. deviation | | |
| Bond lengths (Å) | 0.02 | |
| Bond angles (°) | 2.39 | |
| **Validation** | | |
| MolProbity score | 2.11 | |
| Clashscore | 6.40 | |
| Rotamer outliers (%) | 6.00 | |
| Cβ outliers (%) | 0.16 | |
| CaBLAM outliers (%) | 0.94 | |
| Ramachandran plot | | |
| Favored (%) | 97.06 | |
| Allowed (%) | 2.78 | |
| Outliers (%) | 0.15 | |

**Table S2.**

**Orthologous subunits in Mrp antiporters, MBH, MBS and respiratory complex I**

| Mrp | MBH | MBS | Complex I (human) | Complex I (bacterial) |
|---|---|---|---|---|
| MrpA (TM 1-14) | - | MbsH’ | ND5 | Nqo12 |
| MrpA (TM 15,16 + lateral helix) | MbhI (C-terminal) | | | |
| MrpA (TM 17-19) | MbhD | MbsD | ND6 | Nqo10 |
| MrpA (TM 20-21) | MbhE | MbsE | | |
| MrpB | MbhF | | - | - |
| MrpC | MbhG | MbsG | ND4L | Nqo11 |
| MrpD | MbhH | MbsH | ND2 / ND4 | Nqo13 / Nqo14 |
| MrpE | MbhA | MbsA | - | - |
| MrpF | MbhB | MbsB | - | - |
| MrpG | MbhC | MbsC | - | - |

**Table S3. Identifier for water molecules in the hydrophobic transmembrane region.**

| **water** | water PDB file | **water** | water PDB file |
|---|---|---|---|
| **1** | 255 | **40** | 315 |
| **2** | 254 | **41** | 346 |
| **3** | 344 | **42** | 94 |
| **4** | 212 | **43** | 95 |
| **5** | 211 | **44** | 97 |
| **6** | 29 | **45** | 72 |
| **7** | 30 | **46** | 71 |
| **8** | 31 | **47** | 73 |
| **9** | 32 | **48** | 13 |
| **10** | 38 | **49** | 12 |
| **11** | 279 | **50** | 11 |
| **12** | 280 | **51** | 15 |
| **13** | 33 | **52** | 16 |
| **14** | 27 | **53** | 17 |
| **15** | 28 | **54** | 18 |
| **16** | 26 | **55** | 21 |
| **17** | 25 | **56** | 19 |
| **18** | 24 | **57** | 20 |
| **19** | 273 | **58** | 200 |
| **20** | 364 | **59** | 126 |
| **21** | 1 | **60** | 131 |
| **22** | 274 | **61** | 127 |
| **23** | 3 | **62** | 128 |
| **24** | 6 | **63** | 129 |
| **25** | 2 | **64** | 130 |
| **26** | 5 | **65** | 132 |
| **27** | 281 | **66** | 133 |
| **28** | 350 | **67** | 134 |
| **29** | 4 | **68** | 152 |
| **30** | 220 | **69** | 153 |
| **31** | 7 | **70** | 268 |
| **32** | 8 | **71** | 269 |
| **33** | 343 | **72** | 169 |
| **34** | 283 | **73** | 367 |
| **35** | 9 | **74** | 361 |
| **36** | 14 | **75** | 165 |
| **37** | 10 | **76** | 166 |
| **38** | 353 | **77** | 366 |
| **39** | 93 | **78** | 170 |

**Table S4 Conservation of residues in putative ion translocation pathways**

| Mrp-Subunit | Residue[a] | Conservation in Mrp [b] | Conservation in Complex I [b] (Substitutions) | | Mutants with no or minor impact on activity [c] | Mutants with negative impact on activity [c] |
|---|---|---|---|---|---|---|
| | | | ND2 | ND4 | | |
| MrpD | Glu137 | strictly | strictly | strictly | E137D[2,5] | E137A[2,5,9]; E137Q[2,5] |
| MrpD | Met139 | moderately (Met or Leu) | No | No | - | - |
| MrpD | Ser143 | strictly | No | No | - | - |
| MrpD | Tyr162 | highly | strictly | No | - | - |
| MrpD | Gln166 | moderately | No | No (strictly Thr) | - | - |
| MrpD | Ser170 | moderately | moderately | highly | - | - |
| MrpD | Leu172 | No | No | No | - | - |
| MrpD | Met218 | No | No (moderately Val or Leu) | No (moderately Phe or Met) | - | - |
| MrpD | Lys219 | strictly | strictly | strictly | - | K219A[2]; K220A[9] |
| MrpD | Tyr233 | strictly | moderately | No (highly His) | - | - |
| MrpD | Thr249 | highly | No | No (highly Leu) | - | - |
| MrpD | Lys250 | strictly | strictly | strictly | - | K251A[9] |
| MrpD | Asp295 | No | No | highly | - | - |
| MrpD | Lys297 | moderately (Lys or Arg) | highly | strictly | - | - |
| MrpD | His303 | highly* (Asn or His) | No (highly Ser) | No (highly Ser) | - | - |
| MrpD | Ser306 | moderately* (Ser or No) | No | moderately (Ala or Ser) | - | - |
| MrpD | Gln307 | moderately* (Gln or small) | No (highly His or Asn) | No (highly His) | - | - |
| MrpD | Tyr328 | moderately (Phe or Tyr) | No | No | - | - |
| MrpD | Tyr329 | highly | moderately | highly | - | - |
| MrpD | His332 | highly | No | No | - | - |
| MrpD | His333 | highly* (Asp or His) | No (highly Tyr) | highly | - | - |
| MrpD | Lys337 | highly* | No (moderately Asn or Thr) | No (moderately Ser or Thr) | - | - |
| MrpD | Phe341 | highly (Phe or Tyr) | highly | highly | - | F341A[3] |
| MrpD | Lys392 | strictly | strictly | No (strictly Glu) | - | K392A[9] |
| MrpD | Phe393 | No | No | No | - | - |
| MrpD | Met423 | No | No (moderately Ile, Val or Leu) | No (moderately Leu or Ile) | - | - |

| Mrp-Subunit | Residue[a] | Conservation in Mrp [b] | Conservation in Complex I [b] (Substitutions) | Mutants with no or minor impact on activity [c] | Mutants with negative impact on activity [c] |
|---|---|---|---|---|---|
| MrpA | Tyr101 | strictly | highly | - | - |
| MrpA | Met118 | No | No | - | - |
| MrpA | Phe119 | strictly | highly | - | - |
| MrpA | Tyr136 | No | No (highly Phe) | Y136A[2] | - |
| MrpA | Trp139 | strictly | strictly | - | - |
| MrpA | Glu140 | strictly | strictly | E113Q[6] | E113Q[5]; E132A[9]; E140A[2] |
| MrpA | Ser143 | highly | No (highly Gly) | - | - |
| MrpA | Ser146 | strictly | highly | - | - |
| MrpA | Ser147 | No | No | - | - |
| MrpA | Met167 | No | No | - | - |
| MrpA | Thr170 | strictly | No (strictly Asn) | - | - |
| MrpA | Thr222 | highly | No (highly Gly) | - | - |
| MrpA | Lys223 | strictly | strictly | K196A[6] | K223A[2]; E213A[9] |
| MrpA | Pro240 | highly | strictly | - | - |
| MrpA | Thr241 | highly | strictly | - | - |
| MrpA | Pro242 | strictly | strictly | - | - |
| MrpA | Val243 | highly | highly | - | - |
| MrpA | Ser244 | strictly | strictly | - | - |
| MrpA | Ala245 | highly | highly | - | - |
| MrpA | Tyr246 | highly | No (highly Leu) | - | - |
| MrpA | Leu247 | strictly | moderately | - | - |
| MrpA | His248 | strictly | strictly | H221A[6] | H248A[10] |
| MrpA | Ser249 | highly | No (highly Ala) | - | - |
| MrpA | Ala250 | strictly | moderately | - | ΔA240[9] |
| MrpA | Thr251 | highly | highly | T224A[6] | - |
| MrpA | Met252 | highly | highly | M225I[6] | - |
| MrpA | Lys254 | strictly | No (moderately Thr) | K244A[9] | - |
| MrpA | Asp297 | highly | strictly | - | - |
| MrpA | Lys299 | strictly | strictly | K299A[2] | - |
| MrpA | Thr306 | strictly | strictly | - | - |
| MrpA | Ser308 | highly | strictly | - | - |
| MrpA | Gln309 | moderately (Gln or His) | highly | - | - |
| MrpA | His345 | highly | strictly | H345A[2] | - |
| MrpA | Leu346 | moderately (Leu or Ile) | moderately (Leu or Val) | - | - |
| MrpA | His349 | strictly | strictly | - | - |
| MrpA | Lys353 | strictly | strictly | K329A[9] | - |
| MrpA | Ser407 | highly | highly | - | - |
| MrpA | Lys408 | strictly | strictly | - | K384A[9] |
| MrpA | Glu409 | strictly | strictly | - | E385A[9] |
| MrpA | Thr413 | No | No | - | - |
| MrpA | Thr444 | moderately | highly | - | - |
| MrpA | Tyr447 | highly | strictly | - | - |
| MrpA | Tyr525 | No | No | - | - |

| Mrp-Subunit | Residue[a] | Conservation in Mrp [b] | Conservation in Complex I [b] (Substitutions) | Mutants with no or minor impact on activity [c] | Mutants with negative impact on activity [c] |
|---|---|---|---|---|---|
| MrpA | His528 | highly | No | - | - |
| MrpA | Asn531 | No | No | - | - |
| MrpA | Glu533 | No | No | - | - |
| MrpA | Thr537 | moderately (Thr or Ser) | No (moderately Lys or Glu) | - | - |
| MrpA | Ile650 | No | No | - | - |
| MrpA | Ala656 | No | No | - | - |
| MrpA | Val657 | No | No | - | - |
| MrpA | Val660 | No | No | - | - |
| MrpA | Asp678 | strictly | No | D647A[9] | - |
| MrpA | Thr682 | strictly | No | T683A[3] | - |
| MrpA | Gln683 | strictly | No | - | - |
| MrpA | Val686 | moderately | No | - | - |
| MrpA | Glu687 | strictly | No | E657D[5] | E687A[3]; E656A[9] |
| MrpA | Thr688 | No | No | - | - |
| MrpA | Thr690 | moderately (Ser or Thr) | No | - | - |
| MrpA | Val691 | No | No | - | - |
| MrpA | Leu694 | No | No | - | - |
| MrpA | Leu704 | No | No | - | - |
| MrpA | Glu707 | No | No | - | - |
| MrpA | Asn766 | strictly | No | - | - |
| MrpA | Asp771 | highly | No | D736A[9] | - |
| MrpA | Asp776 | strictly | No | D743E[5] | D743N[5]; D741A[9] |
| MrpA | Thr777 | strictly | No | - | - |
| MrpA | Glu780 | strictly | No | - | E747Q[4,5]; E780A[3]; E747A[4]; E745A[9]; E747D[4,5] |
| MrpB | His34 | strictly | - | H34A[2] | - |
| MrpB | Pro37 | strictly | - | - | P37G[2] |
| MrpB | Gly38 | strictly | - | - | - |
| MrpB | Gly39 | strictly | - | - | - |
| MrpB | Gly40 | strictly | - | - | - |
| MrpB | Phe41 | strictly | - | - | F41A[2] |
| MrpB | Glu111 | No | - | - | - |
| MrpB | Glu113 | No | - | - | - |
| MrpB | Asp121 | highly | - | D121E[5] | D121A[5]; D121N[5] |
| MrpC | Leu26 | No | No | - | - |
| MrpC | Arg27 | moderately (Arg, Gln, or Lys) | No | - | - |
| MrpC | Val30 | No | No | - | - |
| MrpC | Gly31 | highly | No | - | - |
| MrpC | Ser36 | moderately | No | - | - |
| MrpC | His37 | moderately (His, Tyr or Asn) | No | - | - |
| MrpC | His40 | moderately (His or Asn) | No | - | - |

| Mrp-Subunit | Residue[a] | Conservation in Mrp [b] | Conservation in Complex I [b] (Substitutions) | Mutants with no or minor impact on activity [c] | Mutants with negative impact on activity [c] |
|---|---|---|---|---|---|
| MrpC | Leu74 | strictly | No | - | - |
| MrpC | Thr75 | strictly | No (highly Ala) | - | T75A[2] |
| MrpC | Ala76 | moderately (Ala or Ser) | moderately | - | - |
| MrpC | Ile77 | strictly | No | - | I76F[7] |
| MrpC | Val78 | strictly | No (highly Glu) | - | - |
| MrpC | Ile79 | strictly | No | - | - |
| MrpC | Ser80 | No | No | - | - |
| MrpC | Thr84 | moderately | No | - | - |
| MrpC | Ser85 | No | No | - | - |
| MrpE | Thr113 | highly | - | - | T113A[1,2]; T113Y[2] |
| MrpE | Thr116 | moderately (Thr or Ser) | - | - | - |
| MrpE | Met119 | No | - | - | - |
| MrpE | His131 | strictly | - | H131A[1] | - |
| MrpF | Met13 | No | - | - | - |
| MrpF | Ile34 | Moderately (Val or Ile) | - | - | - |
| MrpF | Asp38 | highly | - | D36A/F40D[9]; D38N[5]; D36A/I33D[9] | D38A[5]; D36A[9]; D35L[8]; D36L[9]; D36N[9]; D38E[5] |
| MrpF | Thr39 | No | - | - | - |
| MrpF | Asn43 | No | - | - | - |
| MrpF | Ser68 | No | - | | |
| MrpF | Ser75 | Moderately Thr | - | - | - |
| MrpG | Ser19 | No | - | - | - |
| MrpG | Ser23 | No | - | - | - |
| MrpG | His37 | Moderately | - | - | - |
| MrpG | Thr40 | moderately (Thr or Ser) | - | - | - |
| MrpG | Thr44 | moderately (Thr or Ser) | - | - | - |
| MrpG | Phe75 | highly | - | - | - |
| MrpG | Thr79 | highly | - | - | - |

[a] Residues that show multiple conformations in our structure are underlined.

[b] The conservation is given in four categories based on alignments of 1200-2000 non-redundant sequences: strict (>99.5%); high (>90%); moderate (>80% or >90% in case residues with similar properties are conserved at this position) or not conserved (deviating from BpMrp).

[c] Mutations are given in short form with the sequence numbering from the original source and the source given in superscript: [1] (*24*); [2] (*23*); [3] (*25*); [4] (*31*); [5] (*29*); [6] (*65*); [7] (*66*); [8] (*14*); [9] (*15*); [10] This work

* Refers to group 1 operons (separate MrpA and MrpB) only. See Figure S7

**Table S5.** Model systems and simulation time scales.

| System | Conformation# | Charge state | Simulation length |
|---|---|---|---|
| SA1 | *A* | Standard | 3 x 500 ns |
| SB1 | *B* | Standard | 3 x 500 ns |
| PA1 | *A* | Propka-based | 3 x 500 ns |
| PB1 | *B* | Propka-based | 3 x 500 ns |
| PBE | *B* | Propka-based except $H248^{MrpA}$ HSE* | 3 x 500 ns |
| PBP | *B* | Propka-based except $H248^{MrpA}$ HSP** | 3 x 500 ns |
| SMA1 | *A* | Standard, H248A | 500 ns |
| SMB1 | *B* | Standard, H248A | 500 ns |
| PMA1 | *A* | Propka, H248A | 500 ns |
| PMB1 | *B* | Propka, H248A | 500 ns |
| SNA1 | *A* | Standard, sodium ion modelled near anionic $D38^{MrpF}$ | 800 ns |
| SNA2 | *A* | Standard, snapshot from SNA1, but $D38^{MrpF}$ neutral | 160 ns |
| SNA3 | *A* | Standard, snapshot from SNA2, but $E687^{MrpA}$ neutral | 800 ns |
| SNA4 | *A* | Standard, snapshot from SA1 but sodium ion modelled in the hydrated path towards $E687^{MrpA}$ | 100 ns, 13 ns |
| SNA5 | *A* | Standard, snapshot from SA1, except $E687^{MrpA}$, $D771^{MrpA}$, $D678^{MrpA}$, $D121^{MrpB}$, $E113^{MrpB}$ and $D38^{MrpF}$ neutral. Sodium ion modelled in between $E687^{MrpA}$ and $H37^{MrpC}$. | 110 ns |
| SNA6 | *A* | Standard, snapshot from SNA5 but $H37^{MrpC}$ and $H40^{MrpC}$ doubly-protonated. | 280 ns, 110 ns |
| SNA7 | *A* | Snapshot from SNA6, but $E137^{MrpD}$ neutral. | 2 x 110 ns |

### Alternative location A and B, as defined in PDB file.

* HSE means neutral histidine with ε nitrogen protonated

** HSP means that histidine is doubly-protonated

**Table S6.** Protonation states of amino acids in *A* and *B* conformations based on pKa calculations (see methods)

| | | Conformation *A* | | Conformation *B* | |
|---|---|---|---|---|---|
| **Subunit** | **Residue** | **pKa** | **Charge state** | **pKa** | **Charge state** |
| MrpA | Asp678 | 7.24 | 0 | 7.36 | 0 |
| MrpA | Asp771 | 7.65 | 0 | 7.65 | 0 |
| MrpA | Glu409 | 7.20 | 0 | 6.79 | -1 |
| MrpA | Glu687 | 8.42 | 0 | 7.75 | 0 |
| MrpA | Glu780 | 8.54 | 0 | 9.52 | 0 |
| MrpA | His470 | 7.17 | +1 | 7.17 | +1 |
| MrpA | Lys223 | 6.94 | 0 | 6.65 | 0 |
| MrpA | Lys254 | 5.85 | 0 | 5.90 | 0 |
| MrpA | Lys299 | 6.86 | 0 | 6.84 | 0 |
| MrpA | Lys353 | 6.40 | 0 | 6.43 | 0 |
| MrpA | Lys408 | 7.94 | +1 | 6.37 | 0 |
| MrpB | Asp121 | 7.34 | 0 | 7.35 | 0 |
| MrpD | Lys250 | 6.31 | 0 | 6.31 | 0 |
| MrpD | Lys337 | 5.61 | 0 | 5.61 | 0 |
| MrpF | Asp38 | 6.70 | -1 | 7.03 | 0 |
